# Genomic structure, evolutionary origins, and reproductive function of a large amplified intrinsically disordered protein gene on the X chromosome (*Laidx*) in mice

**DOI:** 10.1101/2020.01.08.898924

**Authors:** Martin F. Arlt, Michele A. Brogley, Evan R. Stark-Dykema, Yueh-Chiang Hu, Jacob L. Mueller

## Abstract

Mouse sex chromosomes are enriched for co-amplified gene families, present in tens to hundreds of copies. Co-amplification of *Slx*/*Slxl1* on the X chromosome and *Sly* on the Y chromosome are involved in dose-dependent genetic conflict, however the role of other co-amplified genes remains poorly understood. Here we demonstrate that the co-amplified gene family *Srsx*, along with two additional partial gene annotations, is actually part of a larger transcription unit, which we name *Laidx*. *Laidx* is harbored in a 229 kb amplicon that represents the ancestral state as compared to a 525 kb Y-amplicon containing the rearranged *Laidy*. *Laidx* contains a 25,011 nucleotide open reading frame, predominantly expressed in round spermatids, predicted to encode an 871 kDa protein. *Laidx* has orthologous copies with the rat and also the 825-MY diverged parasitic Chinese liver fluke, *Clonorchis sinensis*, the likely result of a horizontal gene transfer of rodent *Laidx* to an ancestor of the liver fluke. To assess the male reproductive functions of *Laidx*, we generated mice carrying a multi-megabase deletion of the *Laidx*-ampliconic region. *Laidx*-deficient male mice do not show detectable reproductive defects in fertility, fecundity, testis histology, and offspring sex ratio. We speculate that *Laidx* and *Laidy* represent a now inactive X versus Y chromosome conflict that occurred in an ancestor of present day mice.

## Introduction

The mouse has highly co-amplified gene families on the X and Y chromosomes [1]. These co-amplified X- (*Slx*, *Slxl1*, *Sstx*, *Srsx, Astx*) and Y-linked (*Sly*, *Ssty1*, *Ssty2*, *Srsy, Asty*) gene families are predominantly expressed in post-meiotic testicular germ cells, suggesting an important function in reproduction and testicular germ cell development [1–6]. The co-amplification of gene families on the X and Y chromosomes is thought to have arisen because of meiotic drive, the unequal transmission of an allele to the next generation [1, 7, 8]. *Slx*, *Slxl1*, and *Sly* are meiotic drivers, where increases in gene expression generates a competitive advantage of X- or Y-bearing sperm [7, 9]. The *Slx*/*Slxl1* gene family is also required for male fertility, highlighting how a co-amplified gene family can become essential for male fertility [7, 10]. However, little is known about the biological functions or evolutionary origins of other X- and Y-linked co-amplified gene families.

We chose to explore the evolutionary origins and reproductive function of *Srsx* for multiple reasons. First, *Srsx* and *Srsy* share the highest level of nucleotide identity (~95%) of the X- and Y-linked co-amplified gene families in mice [1]. Second, it is unclear if *Srsx* encodes a protein. Third, while we know *Srsx* is present in ~14 copies within a ~2 Mb amplicon array on the X chromosome [2, 11], the evolutionary origins of *Srsx* and the amplicon containing it are not well-defined. Finally, similar to *Slx*/*Slxl1*/*Sly*, *Srsx* is predominantly expressed in round spermatids [1, 2], suggesting a potential role in meiotic drive and male fertility.

## Results

### The Srsx-amplicon is rearranged on the Y chromosome

To define the genomic structure of a single amplicon containing *Srsx*, we generated a high-quality assembly of BAC RP23-106J7 using PacBio sequencing. Comparison of the 209 kb BAC sequence to the mouse reference genome (mm10) revealed the amplicon size is 20 kb larger than the BAC (Figure 1a). We used this 229 kb representative amplicon (chrX:123,326,277-123,555,768; mm10) for all subsequent analyses (Figure 1a, Supplemental Figure 1). The representative *Srsx*-amplicon sequence shares 99.1% sequence identity with another *Srsx*-amplicon (chrX:123,104,838-123,326,276; mm10), differing primarily by L1 and ERVK transposable element content (Supplemental Figure 2). There are truncated forms of the 229 kb amplicon ranging from 120-130 kb in length, which are arranged in tandem in the opposite (palindromic) orientation of the full-length amplicons (Figure 1a, Supplemental Figure 1). Within both the full-length and truncated *Srsx*-amplicons are two internal tandem repeats, one that is 1149 bps, repeated ~12 times and shares 93% sequence identity between repeats, while the other is 381 bps, repeated four times, and shares 96% sequence identity between repeats (Figure 1b). We expect the entire *Srsx* ampliconic region is comprised primarily of these full-length and truncated amplicons, but because there are multiple gaps across the region the genomic architecture of the entire region is unresolved (Figure 1a, Supplemental Figure 1).

**Figure 1.**
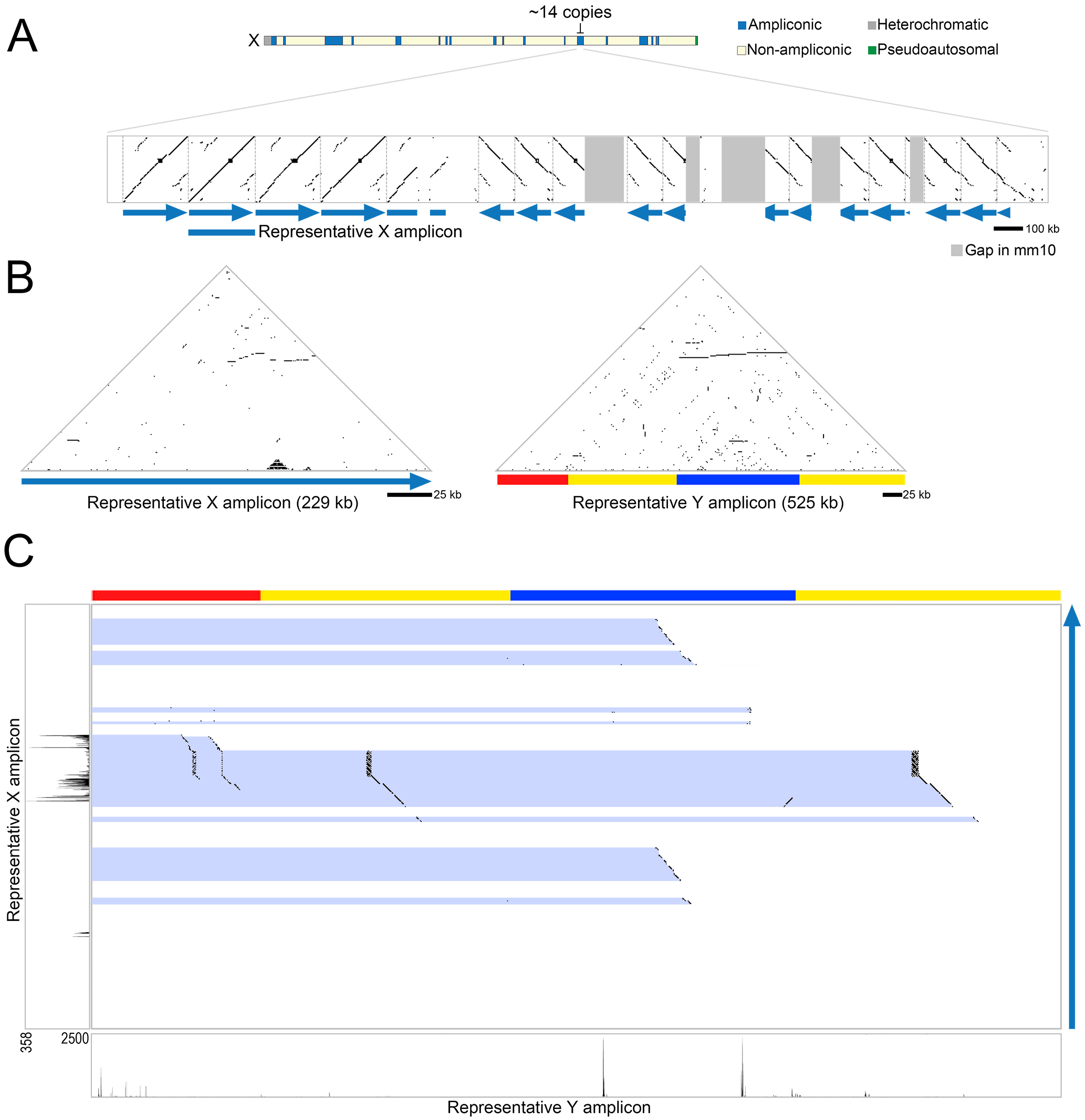
Amplicons containing *Srsx* and*Srsy* are present in multiple copies within ampliconic regions of the mouse X and Y chromosomes. (A) Dot plot comparing the representative *Srsx*-containing amplicon to the entire ampliconic region (chrX:123,050,000-126,250,000; mm10). Each dot represents 100% sequence identity in a 100 bp window. Blue arrows indicate position and orientation of *Srsx*-containing amplicons. The representative sequence used for subsequent analyses is indicated by a blue bar. Vertical dotted gray lines mark the boundaries of each amplicon. Dark gray bars mark gaps in the mm10 reference genome sequence. (B) Self-symmetry triangular dot plots of X- and Y-amplicons are shown with each amplicon compared to itself. Each dot represents a perfect match of 50 nucleotides. Horizontal lines indicate tandemly-arrayed amplicons. The chromosomal regions shown are chrX:123,326,277-123,555,768 and chrY:49,567,447-50,092,166 (mm10). (C) Dot plots of DNA sequence identity between the X- and Y-amplicons from (B), on the Y and X axes, respectively. Each dot represents 100% sequence identity in a 25 bp window. Blue highlights indicate regions of sequence identity between the two sequences. The red, yellow, and blue Y-amplicon subunits are shown at top. Testis RNA-seq reads for each region are illustrated along the respective axes. The positions of *Sly* and *Ssty1/2* have been previously mapped [1] and are excluded from the Y-amplicon for simplicity.

Defining the *Srsx*-amplicon allows us to perform a more accurate comparison with *Srsy*-amplicons. *Srsy* is within a 525 kb amplicon on the Y chromosome also containing *Sly, Ssty1* and *Ssty2* [1]. The 525 kb representative *Srsy*-amplicon consists of three subunits, labeled red, yellow, and blue, with the yellow subunit duplicated within the amplicon (Figure 1b) [1]. Pairwise sequence comparisons between *Srsx-* and *Srsy-* amplicons (Figure 1c), reveal previously observed regions with high levels of nucleotide identity [1]. We additionally find the *Srsx*-amplicon is not contained in its entirety, nor contiguously, within the larger *Srsy*-amplicon. For example, a 34.2 kb region of the *Srsx*-amplicon is represented once in each yellow repeat, as well as twice in degenerated form in the red repeat, while a different part of the *Srsx*-amplicon is duplicated in the blue repeat sequence (Figure 1c). Based on these observations, we speculate the *Srsx*-amplicon represents the ancestral state of a common sequence shared on the X and Y chromosomes.

### Laidx is a large gene in the Srsx-amplicon

We examined how differences between *Srsx*- and *Srsy*-amplicon affect transcription in the testis. Mapping of previously published total RNA-seq sequences [12] from round spermatids to the *Srsx*- and *Srsy*-amplicons reveals a single, long transcription unit in the *Srsx*-amplicon. Sequences homologous to the long *Srsx*-amplicon transcription unit are rearranged within the *Srsy*-amplicon (Figure 1c), suggesting the *Srsy*-amplicon lacks the ability to generate a contiguous transcript homologous to a transcript from the *Srsx*-amplicon. Instead, the rearranged *Srsy*-amplicon sequences produce several, separate transcripts with homology to small segments of the X-amplicon transcript, including *Srsy*, *Asty*, and *Gm28689*. These Y-specific transcripts are detected at low levels (FPKM = 0.03 – 2.13; Supplemental Figure 3). The presence of a single, long, transcription unit within the *Srsx*-amplicon, as compared to fragmented transcripts on the *Srsy*-amplicon further supports the *Srsx*-amplicon as the ancestral state.

We further characterized the large transcription unit within the *Srsx*-amplicon to determine whether it encodes a protein. The large transcription unit spans 35.6 kb and encodes a 28.2 kb mature transcript comprising nine exons (Figure 2a). Consistent with a single transcription start site, reanalysis of ChIP-seq data [13] reveals a small enrichment of RNA polymerase II and a broad enrichment of H3K4me3 overlapping the transcription start site (Figure 2a). This long transcription unit encompasses *Srsx* and two other partial gene annotations also co-amplified on the X and Y chromosomes, *Astx2* and *Gm17412* [6]. There is no enrichment of H3K4me3 at the annotated start sites of *Srsx*, *Astx2*, and *Gm17412*, suggesting they are not independently transcribed in round spermatids. This long transcription unit contains nine exons with exon 1 comprising >93% of the predicted transcript and the entirety of the predicted open reading frame. This long transcription unit is detectable in round spermatids of the testis (Figure 2b, Supplemental Figure 4) and undetectable across several somatic tissues (Supplemental Figure 4), germinal vesicles or oocytes (Figure 2b), consistent with previous studies [6]. Altogether, we find three partially-annotated genes (*Srsx*, *Astx*, and *Gm17412*) are contained within a single transcriptional unit expressed in round spermatids.

**Figure 2.**
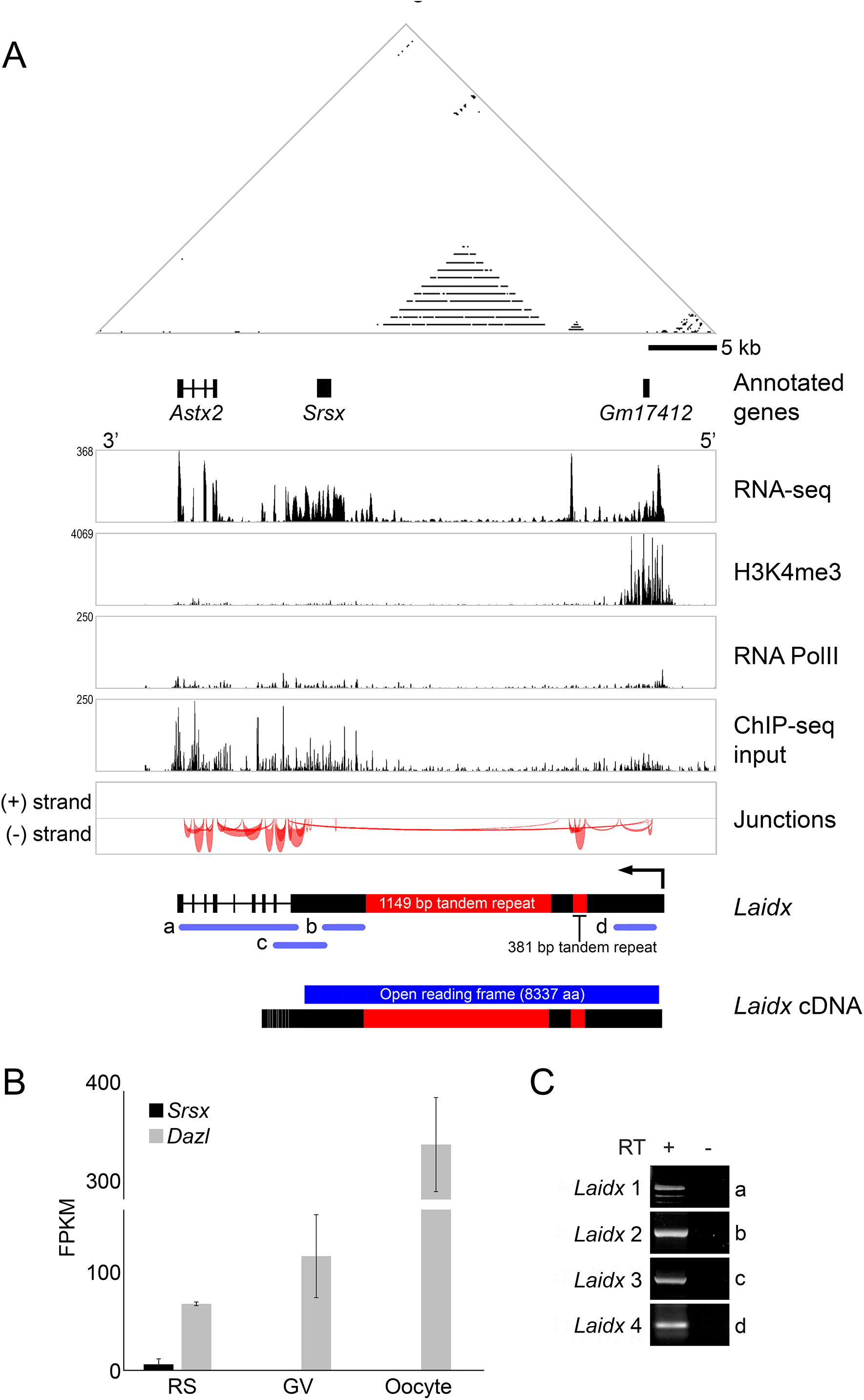
A large transcription unit, *Laidx*, encompasses three partially annotated genes, including *Srsx*. (A) Self-symmetry triangular dot plot of a 45.5 kb region encompassing the large transcription unit. The coordinates of the chromosomal regions shown are chrX:123,443,060-123,488,539 (mm10). Positions of three partially annotated genes (*Astx2*, *Srsx*, and *Gm17412*) are indicated. Below the dotplot and gene annotations are aligned reads from RNA-seq performed on testis, showing transcription extending from upstream of *Gm17412* to the end of *Astx2*. In addition, ChIP-seq revealed modest enrichment of RNA polymerase II along the transcription unit and a small amount of enrichment at the transcription start site, along with broad enrichment of H3K4me3, a modification associated with active promoters. RNA-seq and ChIP-seq alignments were performed on repeat masked sequence [22]. “Junctions” illustrates predicted splice sites based on RNA-seq. (B) Quantitation of RNA-seq data from round spermatids (RS), germinal vesicles (GV), and oocytes demonstrating transcription in the male but not female germline. *Dazl*, a gene expressed in both male and female germlines, is used as a control. FPKM, Fragments Per Kilobase per Million reads. (C) RT-PCR on RT (+) and no RT controls (−) performed on RNA isolated from adult testis.

To validate this novel long gene, we performed RT-PCR with primers specific to different regions of the putative transcription unit (Figures 2a and c, Supplemental Figure 5). We used sequences spanning the intron-exon junctions predicted by Cufflinks [14] to design primers that amplify products spanning multiple exons. These RT-PCR products confirm the expression of a single large transcription unit in testis. While it is not clear if this transcript produces a protein, there is an open reading frame of 25,011 base pairs encoding a large predicted protein of 8337 amino acids (871 kDa). This protein has no known functional motifs and is predicted to be an intrinsically disordered protein (Supplemental Figure 6). We name this new gene, *Laidx* (<u>L</u>arge <u>a</u>mplified <u>i</u>ntrinsically <u>d</u>isordered protein gene encoded on the <u>X</u>). Based on genomic rearrangements and RNA-seq data (Figure 1c), we consider *Laidy* to be pseudogenized in present day mice. The remainder of this study will focus on *Laidx*.

### Laidx migrated between rodents and fluke via horizontal gene transfer

*Laidx* is detectable and potentially amplified on the rat X chromosome, but not detectable in the genomes of guinea pig or deer mouse. We detect 76% nucleotide sequence identity between a rat BAC (CH230-1D6; GenBank Accession AC130042) containing *Laidx* sequence and mouse *Laidx* (Supplemental Figure 7a). While the rat *Laidx* sequence is interrupted by multiple LINE elements, there is a large open reading frame encoding a predicted protein of 1806 amino acids with 43% identity with mouse LAIDX (Supplemental Figure 7b). A comparison of the rat BAC containing *Laidx* BAC to a rat Y BAC (RNAEX-9O8; GenBank Accession AC279156) reveals rat *Laidy* is rearranged, with most of the large open reading frame deleted from the rat Y (Supplemental Figure 8). In both rat and mice, rearrangement of *Laidy* disrupts the coding potential seen in *Laidx*. The conservation of a large open reading frame in mouse and rat suggests *Laidx* is translated.

LAIDX protein shares ~43% sequence identity with a 5280 amino acid hypothetical protein CSKR_14446s (GenBank: RJW68620.1) in the ~825 MY diverged Chinese liver fluke, *Clonorchis sinensis* (Figure 3a). Consistent with the protein similarities, the mouse and *Clonorchis* gene sequences share 69% nucleotide identity across 60% of *Laidx* exon 1 (Figure 3b). Additionally, the 1149 and 381 bp simple tandem repeats in mouse appear single copy in *Clonorchis*.

**Figure 3.**
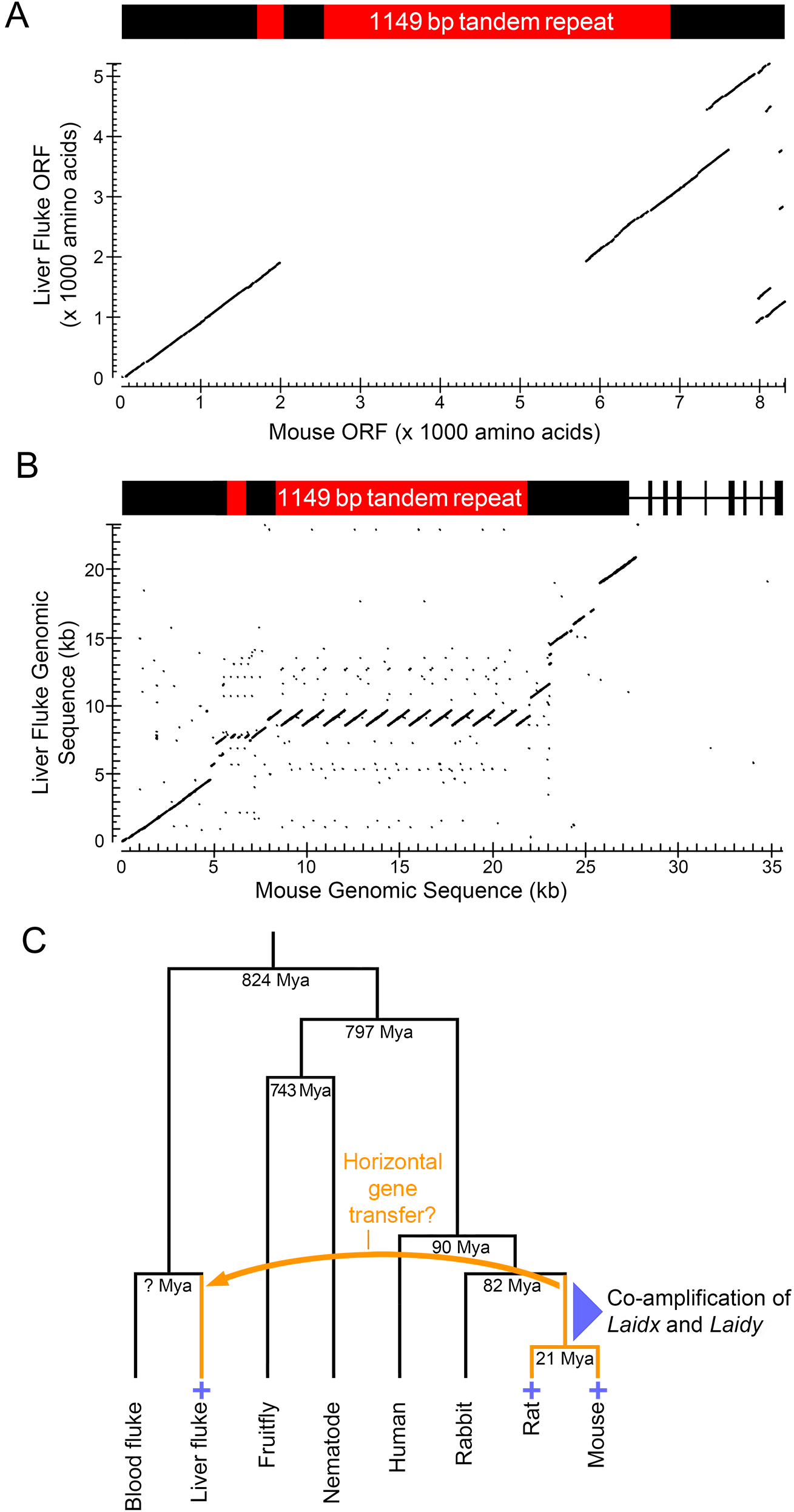
High *Laidx* sequence identity between mouse and Chinese liver fluke suggests a horizontal gene transfer event. (A) Blastp alignment of the 5280 amino acid *Clonorchis* protein CSKR_14446 and the predicted 8337 amino acid LAIDX protein. (B) TBLASTN alignment of *Laidx* with *Clonorchis* genomic sequence encoding CSKR_14446. (C) Evolutionary history of *Laidx*. Species that carry a *Laidx* ortholog are marked with an “+”. An orange arrow and orange branches mark a proposed horizontal gene transfer event that occurred between a mouse/rat ancestor and the liver fluke. Mya, Million years ago.

The conservation of *Laidx* between *Clonorchis*, *Rattus*, and *Mus*, but not in other lineages, including other mammals, insects, and nematodes, suggests *Laidx* moved between rodents and *Clonorchis* via a horizontal gene transfer event in the last 82 million years (Figure 3c). To determine the directionality of the horizontal gene transfer we examined transposable element content in each genomic region. Several rodent-specific ERVK transposable elements are present in the *Clonorchis* sequence encoding CSKR_14446s and non-mammalian transposable elements were not detected in mouse *Laidx*-amplicon. The presence of rodent lineage-specific transposable elements in the *Clonorchis* genome, near this gene, suggests the horizontal gene transfer event occurred with rodent to fluke directionality.

### Laidx deletion and duplication mice do not exhibit overt reproductive defects

To explore the function of *Laidx* in the mouse germline, we generated precise and complete multi-megabase *Laidx* deletions (*Laidx*^−/Y^) and duplications (*Laidx*^Dup/Y^) using CRISPR and Cre/loxP (Figure 4a). Deletions were confirmed by RT-PCR assays specific to three different regions of *Laidx* (Figure 4b). While one assay demonstrated loss of the transcript in the *Laidx*^−/Y^ mice, the other two assays yielded RT-PCR products. Presence of these products could indicate either incomplete deletion or expressed sequences with high sequence identity on the Y chromosome, such as *Srsy* or *Asty*. Sanger sequencing of PCR products from wild-type mice reveals sequence variants between *Laidx* and *Srsy*/*Asty* on the Y chromosome. However, RT-PCR products from the deletion mice contained only Y-specific variants (Figure 4c), supporting that all RT-PCR products are derived from sequences on the Y chromosome, consistent with a complete deletion of the *Laidx*-ampliconic region.

**Figure 4.**
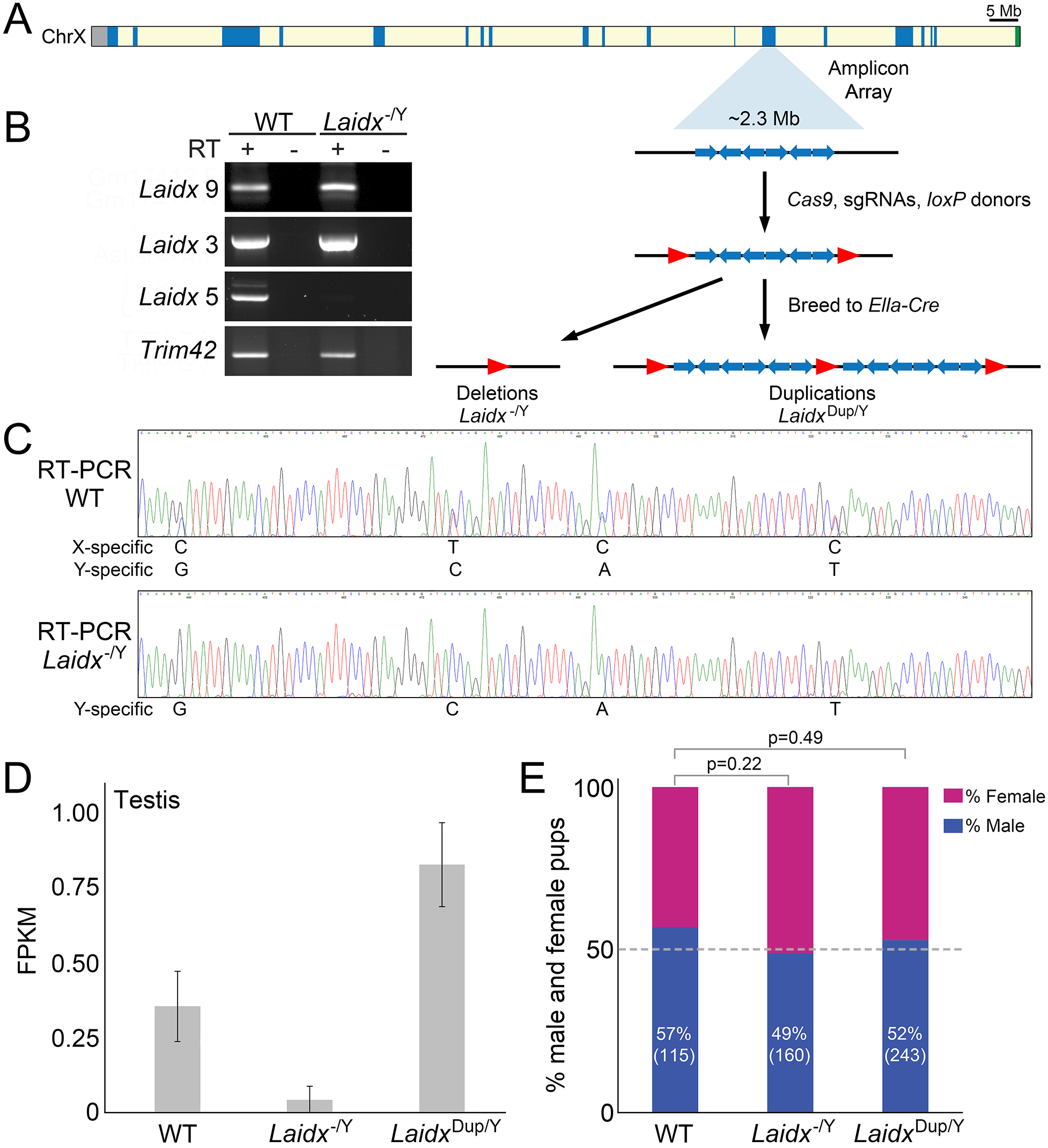
Generation of *Laidx*^−/y^ transgenic mice. (A) Schematic representation of the mouse X chromosome. Ampliconic regions are shown in blue, centromere in gray, and pseudoautosomal region (PAR) in green. The region of the X chromosome carrying the *Laidx*-containing amplicons is expanded to show a representation of the repeat structure (blue arrows). Red arrows denote loxP sites. Mice carrying loxP sites flanking the ampliconic region were mated to *Ella*-Cre mice to generate *Laidx*^−/Y^mice. (B) RT-PCR on RT (+) and no RT controls (−) performed on RNA isolated from adult testis from WT and *Laidx*^−/Y^ mice. *Trim42* is a testis-specific gene used as a positive control. Primer pairs for each assay are indicated (see Supplemental Tables 1 and 2). (C) Sanger sequencing chromatograms from *Laidx* RT-PCR product in WT (top) and *Laidx*^−/Y^ (bottom) mice. The WT product contains multiple sequence variants that are specific to both the X and Y chromosome. The *Laidx*^−/Y^ product contains only variant sequences specific to the Y chromosome. (D) mRNA-seq was performed on testes from WT, *Laidx*^−/Y^, and *Laidx*^Dup/Y^ mice. (E) Males of each genotype were mated against wild-type females and progeny were sex genotyped. The proportion of male offspring is shown as a percentage along with the number of pups screened in parentheses. P-values were calculated using Fisher’s Exact Test.

To further confirm successful deletion and duplication of *Laidx*, mRNA-seq was performed on testes from wild-type, *Laidx*^−/Y^, and *Laidx*^Dup/Y^ mice. Due to the high sequence identity between these regions on the X and Y chromosome, mRNA-seq reads were mapped to a *Laidx* cDNA sequence that is masked across regions with 100% sequence identity between the X and Y chromosome. FPKM of *Laidx*^−/Y^ testes was reduced compared to wild-type mice (FPKM = 0.042 vs 0.354, respectively) supporting deletion of the region (Figure 4d). *Laidx*^Dup/Y^ mice displayed approximately double the level of gene expression (FPKM = 1.81), consistent with a duplication (Figure 4d).

*Laidx*^−/Y^ mice do not display notable reproductive deficits. Testicular morphology, sperm development, and timing of spermatogenic events are not different compared to wild type controls (Supplemental Figure 9). To test the effects of *Laidx* deletion on fertility and fecundity, we bred three *Laidx^−/Y^* males and two wild-type litter mates to wild-type CD1 female mice. *Laidx^−/Y^* male mice have normal fertility and fecundity compared to wild-type (Table 1). *Laidx^−/Y^* males sired 332 pups in 26 litters (mean = 12.8), while wild-type males sired 244 pups in 20 litters (mean = 12.2) (p=0.583). In addition, the litters show no differences in the ratio of male to female pups (Figure 4e). Litters sired by wild-type, *Laidx^−/Y^*, and *Laidx*^Dup/Y^ males are 57% (66/115), 49% (79/160; p=0.22), and 52% (127/243; p=0.49) male, respectively. Compared to wild-type, *Laidx ^−/Y^* males have no change in sperm count or motility, as assessed by the sperm swim-up assay (Table 2). No effect on fertility is observed in *Laidx^−/−^* females.

**Table 1.**
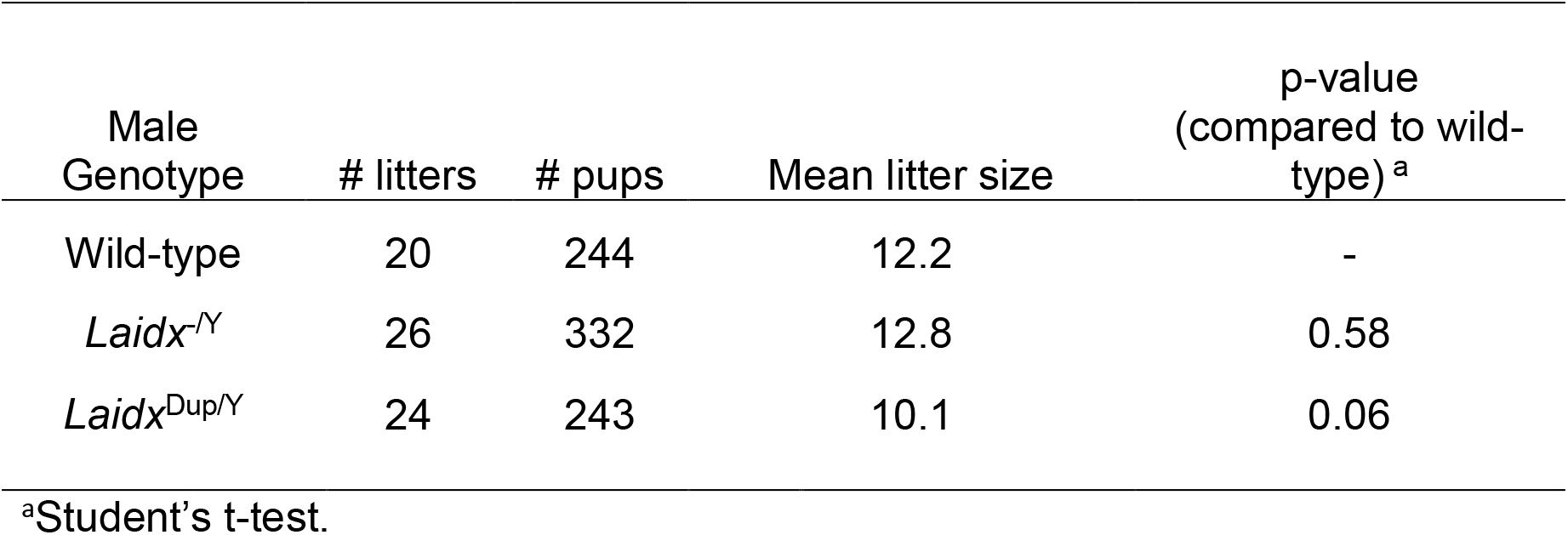
*Srsx*^−/Y^ and *Srsx*^Dup/Y^ male fecundity.

**Table 2.**
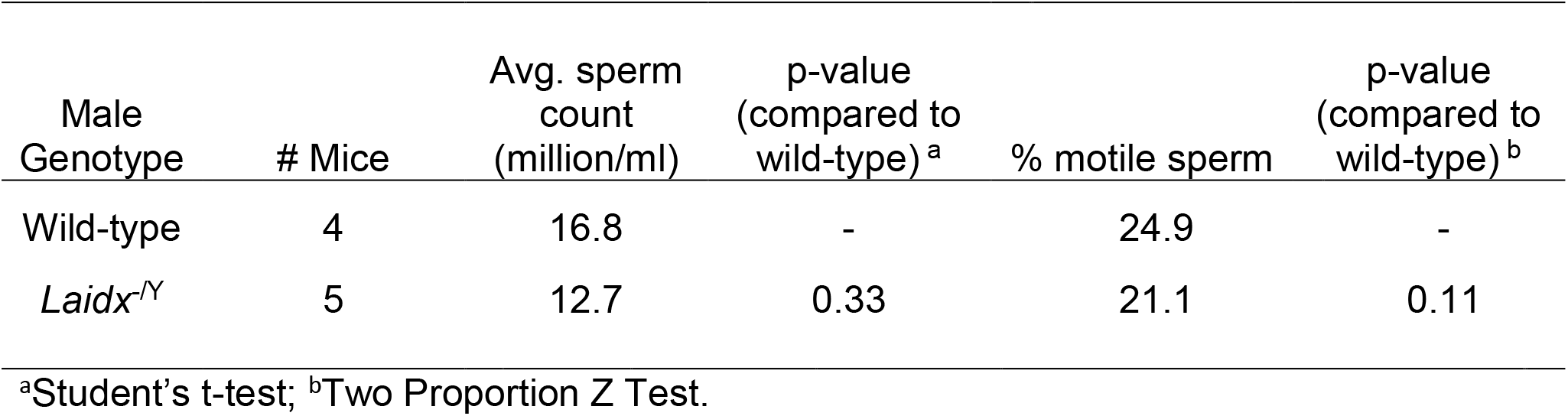
*Srsx*^−/Y^ male sperm count and motility.

## Discussion

We have identified *Laidx*, *a* novel, large, and amplified gene encompassing three previously-annotated genes co-amplified on the mouse X and Y chromosomes. *Laidx* consists of nine exons and encodes an 8337 amino acid (871 kDa) putative protein, which shares sequence similarity with a protein in the Chinese liver fluke, *Clonorchis sinensis*. *Laidx* is predominantly expressed in post-meiotic testicular germ cells, suggesting a role in spermatogenesis and reproduction. However, male mice with *Laidx* deletion and duplication exhibit fertility, fecundity, testis histology, and offspring sex ratio similar to wild-type, indicating the role of *Laidx* may be uncovered under other conditions (e.g. stress, old age). Our resolving the entire gene structure of the large and complex *Laidx* gene sequence combined with the generation of mutant mouse models provides the foundation for future studies exploring the role of *Laidx* in post-meiotic germ cell development.

*Laidx* is present in mouse, rat, and the Chinese liver fluke, *Clonorchis sinensis* and is not detectable in other mammals, suggesting a horizontal gene transfer event. Rat is a definitive host of *Clonorchis* [15], thus providing an opportunity for horizontal gene transfer. While it is difficult to establish horizontal gene transfer conclusively, the presence of rodent-specific ERVK sequences near the *Clonorchis* ortholgous gene suggests *Laidx* was transferred from rodent to the liver fluke. Conservation of a *Laidx* ORF in mouse, rat, and liver fluke along with confirmation of expression in the mouse and rat testis provides additional support for the ancestral state of *Laidx* gene structure and suggests it is important for reproduction. Considering the germ cell expression of *Laidx* in mouse and rat, it will be interesting to examine whether the *Clonorchis* ortholog also functions in germ cells.

The co-amplification of genes on the mouse X and Y chromosomes is thought to have arisen through meiotic conflict, whereby gene duplication confers a competitive advantage in X- or Y-bearing sperm [9]. An example of this phenomenon can be seen in *Sly*, *Slx*, and *Slxl1* [7, 9]. Characterization of *Laidx* reveals that, while there is considerable sequence identity between the *Laidx/y*-amplicons, these chromosomes produce dramatically different transcripts. Given the high similarity of LAIDX to *Clonorchis* CSKR_14446s proteins, we propose that *Laidx* represents the rat/mouse ancestral gene. It is unclear if the massive amplification of *Laidy* is due to functional selection for one of the smaller transcripts, or if it is a passenger resulting from amplification of *Sly*, *Ssty1*, and *Ssty2*, which share the same Y-amplicon [1]. Comparative genomic studies of *Laidx/y* in mammals that predate mouse-rat divergence may provide insights into their role in meiotic drive and the origin of this large predicted protein-coding gene.

## Materials and Methods

### BAC sequencing and assembly

BAC sequencing and assembly was performed as previously described [16]. Briefly, BAC DNA from clones was isolated using a High Pure Plasmid Isolation Kit (Roche Applied Science) following the manufacturer’s instructions. Approximately 1 μg of BAC DNA was sheared using a Covaris g-TUBE. Libraries were processed using the PacBio SMRTbell Template Prep kit following the protocol “Procedure and Checklist—20 kb Template Preparation Using BluePippin Size-Selection System” with the addition of barcoded SMRTbell adaptors. Library size was measured using a FEMTO Pulse. The pooled library was size-selected on a Sage PippinHT with a start value of 12,000 and an end value of 50,000. BACs were sequenced on a PacBio Sequel with version 3.0 chemistry on one SMRT cell and then demultiplexed using LIMA in SMRTlink6.0. Demultiplexed reads were run through the CCS algorithm in SMRTlink6.0. CCS reads were filtered for contaminating *E. coli* reads, and the resulting filtered fasta file was used as input for assembly using Canu v1.8.

### Dot plots

Dot plots showing sequence identity within one sequence and between two sequences were generated using fastdotplot, a custom Perl script that can be found at https://www.pagelab.wi.mit.edu/materials-request. Nucleic acid sequence alignments between *Mus musculus*, *Rattus norvegicus*, and *Clonorchis sinensis* was performed using the blastn algorithm for somewhat similar sequences [17]. Amino acid alignments between *Mus musculus*, *Rattus norvegicus*, and *Clonorchis sinensis* was performed using the blastp algorithm [17].

### mRNA-seq

Total RNA quality was assessed using the 2100 Bioanalyzer (Agilent). 400ng of total RNA (RIN >6) was used to generate polyA-selected libraries with Kapa mRNA HyperPrep kits (Roche) with indexed adaptors. Libraries were assessed on the Tapestation 2200 (Agilent) and quantitated by Kapa qPCR. Pooled libraries were subjected to 50 bp paired-end sequencing according to the manufacturer’s protocol (Illumina NovaSeq6000). Bcl2fastq2 Conversion Software (Illumina) was used to generate de-multiplexed Fastq files.

### RNA-seq and ChIP-seq mapping

RNA-seq and ChIP-seq analyses were conducted by analyzing previously published datasets. Specifically, mouse tissue panel data were analyzed from SRP016501 [18], oocyte data from SRP061454 [19], germinal vesicle data from SRP065256 [20], and sorted round spermatid data from SRP111389 [12]. Rat testis data was analyzed from ERR3417900. Alignments were performed with Tophat, using genomic sequence from the representative X- and Y-amplicons as the reference genome. Due to the ampliconic nature of these sequences --max-multihits were set to 1 and -read-mismatches set to 0; otherwise, standard default parameters were used. We used Cufflinks, with the refFlat RefSeq gene annotation file, to estimate expression levels as fragments per kilobase per millions of mapped fragments (FPKM).

### RNA and RT-PCR

Total RNA was extracted using Trizol (Life Technologies) according to manufacturer’s instructions. Ten μg of total RNA was DNase treated using Turbo DNAse (Life Technologies) and reverse transcribed using Superscript II (Invitrogen) using *Laidx*-specific primers following manufactures instructions. Intron-spanning primers were used to perform RT-PCR on adult testis cDNA preparations for *Laidx* and a round spermatid-specific gene *Trim42* (Supplemental Tables 1 and 2).

### Transgenic lines

To generate mice with multi-megabase deletions of the *Laidx* ampliconic region, loxP sites were sequentially integrated upstream and downstream of the *Laidx* ampliconic region via CRISPR/Cas9. LoxP sites were introduced via cytoplasmic injection of Cas9 mRNA, an sgRNA targeting unique sequence flanking the ampliconic region, and a single stranded oligo donor carrying the loxP sequence (Supplemental Table 3). Cytoplasmic injections were performed on zygotes derived from a F1 (DBA2xC57B6/J) male and a C57B6/J female to ensure all targeted X chromosomes were of C57B6/J origin. Floxed mice were mated against C57B6/J *EIIa-Cre* mice resulting in mice carrying either a *Laidx* region deletion or duplication. Two independent deletion and duplication lines were generated. No differences were observed between independent lines and data were therefore compiled. All deletion or duplication carrying males were derived from heterozygous female mice that had been backcrossed to C57BL/6J males for at least four generations. Mice were genotyped by extracting DNA from a tail biopsy using Viagen DirectPCR lysis reagent using primers that flank loxP sites (Supplemental Table 3).

### Histology

Testes were collected from 2-6 month old mice. The tunica albuginea was nicked and fixed with Bouins Fixative overnight at 4°C. Testes were then washed through a series of ethanol washes (25%, 50%, 75% EtOH) before being stored in 75% EtOH at 25°C. Testes were paraffin embedded and sectioned to 5μm. Sections were stained with Periodic Acid Schiff (PAS) and hematoxylin, visualized using a light microscope, and staged [21]. Specific germ cell populations were identified based upon their location, nuclear size, and nuclear staining pattern [21].

### Fertility, fecundity, and sex ratio distortion assessments

The fecundity of males was assessed by mating at least three deletion and duplication males 2-6 months of age to 2-6 month old CD1 females and monitoring females for copulatory plugs. The fertility of all lines was compared to that of C57B6/J males (wild-type). Offspring sex ratio data were compiled by sex genotyping offspring from the aforementioned crosses as well as pups resulting from males bred with CD1 females. Sex genotyping was done with PCR using primers specific to *Uba1x/y*. Sperm counts were conducted on sperm isolated from the cauda epididymis. Briefly, the cauda epididymis was isolated and nicked three times to allow sperm to swim out. The nicked epididymis was then rotated for 1hr at 37°C in Toyoda Yokoyama Hosi media (TYH). Sperm were fixed in 4% PFA and counted using a hemocytometer. For each genotype, at least three male mice were counted with three technical replicates performed for each mouse and averaged. Testes were collected from 2-6 month old males for all experiments and weighed, along with total body, in order to determine relative testis weight.

### Sperm swim-up assay

Mouse cauda epididymis were dissected from *Laidx*^−/Y^ mice and wild-type litter mate controls. The cauda epididymis was nicked three times to allow sperm to swim out and placed in a 2ml round bottom Eppendorf tube filled with 1.1ml of 37°C Human Tubal Fluid media (Millipore). Sperm were placed in a 37°C incubation chamber and allowed to swim out for 10 minutes before the cauda epididymis was removed. A 30μl aliquot was removed as a pre-swim-up reference. Samples were centrifuged for 5 minutes at 2000 rpm, placed at a 45° angle in a 37°C incubation chamber, and sperm allowed to swim out of the pellet for one hour. The pre- and post-swim-up sperm were counted using a hemocytometer and percent motility calculated. Three technical replicates were performed per mouse.

### Data availability

The complete BAC sequence of RP23-106J7 generated in this study is available from NCBI GenBank (Accession MN842289). All mRNA-seq libraries generated in this study are available at NCBI SRA (PRJNA595938). Supplemental material consisting of three supplemental tables and nine supplemental figures available at figshare.

## Acknowledgements

We would like to acknowledge Dirk de Rooij at Utrecht University for his assistance with histology evaluation, the University of Michigan Histology Core, the University of Michigan Advanced Genomics Core for Sanger and Illumina sequencing and mRNA-seq library preparations, Melanie Sorensen at University of Washington for BAC sequencing, Katherine Stansifer at Ohana Biosciences for the sperm swim-up assay protocol, and the Transgenic Animal and Genome Editing Core at Cincinnati Children's Hospital Medical Center. We thank David C. Page and colleagues at the Whitehead Institute and the Human Genome Sequencing Center at Baylor College of Medicine for the public database deposits of the finished rat Y BAC sequences used in our analyses. We thank David Ginsburg for sharing *EIIa-Cre* mice. We thank Emma Gerlinger and Melody Gorishek for assistance with genotyping. We thank Alyssa Kruger, Callie Swanepoel, and Eden Dulka for helpful comments. This work was supported by National Institutes of Health grant R01HD094736.

## Literature Cited

1. Soh, Y.Q., et al., Sequencing the mouse Y chromosome reveals convergent gene acquisition and amplification on both sex chromosomes. Cell, 2014. 159(4):p. 800–13.

2. Mueller, J.L., et al., The mouse X chromosome is enriched for multicopy testis genes showing postmeiotic expression. Nat Genet, 2008. 40(6): p. 794–9.

3. Szot, M., et al., Does Rbmy have a role in sperm development in mice? Cytogenet Genome Res, 2003. 103(3-4): p. 330–6.

4. Reynard, L.N., J. Cocquet, and P.S. Burgoyne, The multi-copy mouse gene Sycp3-like Y-linked (Sly) encodes an abundant spermatid protein that interacts with a histone acetyltransferase and an acrosomal protein. Biol Reprod, 2009. 81(2): p. 250–7.

5. Comptour, A., et al., SSTY proteins co-localize with the post-meiotic sex chromatin and interact with regulators of its expression. FEBS J, 2014. 281(6): p. 1571–84.

6. Toure, A., et al., Identification of novel Y chromosome encoded transcripts by testis transcriptome analysis of mice with deletions of the Y chromosome long arm. Genome Biol, 2005. 6(12): p. R102.

7. Kruger, A.N., et al., A Neofunctionalized X-Linked Ampliconic Gene Family Is Essential for Male Fertility and Equal Sex Ratio in Mice. Curr Biol, 2019. 29(21): p. 3699–3706 e5.

8. Rathje, C.C., et al., Differential Sperm Motility Mediates the Sex Ratio Drive Shaping Mouse Sex Chromosome Evolution. Curr Biol, 2019. 29(21): p. 3692–3698 e4.

9. Cocquet, J., et al., A genetic basis for a postmeiotic X versus Y chromosome intragenomic conflict in the mouse. PLoS Genet, 2012. 8(9): p. e1002900.

10. Cocquet, J., et al., Deficiency in the multicopy Sycp3-like X-linked genes Slx and Slxl1 causes major defects in spermatid differentiation. Mol Biol Cell, 2010. 21(20): p. 3497–505.

11. Bennett-Baker, P.E. and J.L. Mueller, CRISPR-mediated isolation of specific megabase segments of genomic DNA. Nucleic Acids Res, 2017. 45(19): p. e165.

12. Wichman, L., et al., Dynamic expression of long noncoding RNAs reveals their potential roles in spermatogenesis and fertility. Biol Reprod, 2017. 97(2): p. 313–323.

13. Hammoud, S.S., et al., Chromatin and transcription transitions of mammalian adult germline stem cells and spermatogenesis. Cell Stem Cell, 2014. 15(2): p. 239–53.

14. Trapnell, C., et al., Transcript assembly and quantification by RNA-Seq reveals unannotated transcripts and isoform switching during cell differentiation. Nat Biotechnol, 2010. 28(5): p. 511–5.

15. Chai, J.Y., K. Darwin Murrell, and A.J. Lymbery, Fish-borne parasitic zoonoses:status and issues. Int J Parasitol, 2005. 35(11-12): p. 1233–54.

16. Vollger, M.R., et al., Long-read sequence and assembly of segmental duplications. Nat Methods, 2019. 16(1): p. 88–94.

17. Coordinators, N.R., Database resources of the National Center for Biotechnology Information. Nucleic Acids Res, 2018. 46(D1): p. D8–D13.

18. Merkin, J., et al., Evolutionary dynamics of gene and isoform regulation in Mammalian tissues. Science, 2012. 338(6114): p. 1593–9.

19. Yu, C., et al., Oocyte-expressed yes-associated protein is a key activator of the early zygotic genome in mouse. Cell Res, 2016. 26(3): p. 275–87.

20. Yu, C., et al., BTG4 is a meiotic cell cycle-coupled maternal-zygotic-transition licensing factor in oocytes. Nat Struct Mol Biol, 2016. 23(5): p. 387–94.

21. Ahmed, E.A. and D.G. de Rooij, Staging of mouse seminiferous tubule crosssections. Methods Mol Biol, 2009. 558: p. 263–77.

22. Smit, A., Hubley, R & Green, P. RepeatMasker Open-4.0. 2013-2015. 2015; Available from: <http://www.repeatmasker.org>.

